# Easy to interpret Coordinate Based Meta-Analysis of neuroimaging studies: Analysis of Brain Coordinates (ABC)

**DOI:** 10.1101/2020.12.03.409953

**Authors:** CR Tench, R Tanasescu, CS Constantinescu, DP Auer, WJ Cottam

## Abstract

Functional MRI and voxel-based morphometry (VBM) are common approaches to testing hypotheses by neuroimaging. There is no one optimal generalisable method, and the multiple popular software packages have been shown to produce different results. Furthermore, within each package results may be sensitive to settings, such as smoothing or statistical thresholding, that can be difficult to optimise per hypothesis and can have considerable influence. It is useful, therefore, to have a method to meta-analyse published results from multiple studies that tested a similar hypothesis perhaps using different methods as well as different subjects. Coordinate based meta-analysis (CBMA) offers this using only the commonly reported summary results. It is the aim of CBMA to find those results that indicate replicable effects across studies. However, just like the multiple analysis methods offered for neuroimaging, there are multiple CBMA algorithms each with specific features and empirical parameters/assumptions that directly influence the results. With this in mind a new CBMA method (Analysis of Brain Coordinates; ABC) is presented, with the aim of making interpretation of CBMA results easier by eliminating empirical assumptions where possible and by relating statistical thresholding directly to replication of effect.

## Introduction

Coordinate based meta-analysis (CBMA) is commonly used to estimate effects by analysing multiple independent, but related by a shared hypothesis, neuroimaging studies. It is employed to meta-analyse (amongst others) voxel-based morphometry (VBM) or functional magnetic resonance imaging (fMRI) and uses only reported summary statistics; coordinates and/or statistical effect sizes such as the *t* statistic. These methods are important in neuroimaging if studies use few subjects or employ no principled control of the type 1 error rate so potential for false results is high (Bennett et al. 2009; Kiefer 1953), and when the available analysis options can produce different results even on the same data (Li et al. 2020; Popescu et al. 2016). By analysing multiple studies simultaneously, those results that are replicated across at least some can be identified and are assumed indicative of relevance to the hypothesis. In the absence of whole brain statistical images with which to perform full image based meta-analysis (IBMA), CBMA results can help clarify our understanding, provide testable hypotheses for future prospective studies, or help to test hypothesised effects.

Probably the most popular method of performing CBMA is the activation likelihood estimate (ALE) algorithm(Eickhoff et al. 2009, 2012; Laird et al. 2005; Turkeltaub et al. 2002, 2012). However, there are multiple others (Albajes-Eizagirre et al. 2018; S. G. Costafreda 2012; Sergi G. Costafreda et al. 2009; Radua et al. 2012; C. R. Tench et al. 2017, 2020; Wager et al. 2003, 2007) but using different approaches and assumptions to perform the analysis. Commonly CBMA requires a smoothing kernel to extrapolate the reported coordinates into a voxel-wise analysis. Across the different algorithms the kernel can be Gaussian or spherical, and have full width half max (FWHM), or width, of 10mm to 25mm that can be fixed or dependent on the study sample size. Many algorithms employ randomisation of coordinates into an image space to represent an empirical null hypothesis reflecting the situation of no systematic spatial agreement across studies, and various image spaces are used. Unlike classical meta-analysis, which aims to estimate effects from available evidence, CBMA aims to distinguish replicable effects from those that are study specific. This necessitates a statistical thresholding scheme that can be voxel-wise or cluster-wise, based on family wise error (FWE) or false discovery rate (FDR) (Benjamini and Hochberg 1995), or even uncorrected for multiple voxel-wise tests. Despite the considerable differences between the various algorithms, they have a common aim of producing isolated clusters representing the spatial distribution of reported foci most likely related to the common hypothesis as indicated by consistency of reporting across the studies. It is the anatomical location of these clusters that form the output of CBMA algorithms and the result on which the interpretation and conclusion are based.

The different approaches to CBMA means that choice of method has considerable influence over the results (Ferreira and Busatto 2010) and therefore conclusions. When choosing a CBMA method the direct influence of each specific feature of the algorithm should be understood, but this can be difficult when they are empirical, such as the FWHM, or nonlinear, such as threshold. This article describes analysis of brain coordinates (ABC), which attempts to eliminate empirical assumptions where possible. It is model based, requiring only the human grey-matter (GM), white-matter (WM) or whole-brain (WB) volume as appropriate. Statistical thresholding is aimed at identification of replicable effects and is designed to be simply interpreted as the minimum replicates that the analyst considers meaningful. Clustering is performed on coordinates that survive thresholding using no further empirical parameters. Results are numbered clusters of coordinates that can be interpreted and subjected to further analysis. Software to perform ABC is provided to use freely as part of NeuRoi https://www.nottingham.ac.uk/research/groups/clinicalneurology/neuroi.aspx.

## Methods

### Study Density Model

In ABC the results considered most likely associated with the common hypothesis are those where the reported coordinates from different independent studies fall close together spatially, which is quantified by study density; using study density, rather than coordinate density, is equivalent to the common CBMA tactic of considering studies as random effects (Eickhoff et al. 2009; Wager et al. 2007). The algorithm computes for coordinate *i* the smallest spherical volume, *dV_i_*, encompassing the *k* nearest coordinates (including *i*) from *k* different independent studies, with a minimum allowed volume of *dV_i_* =8mm^3^ imposed in case all fall within a single voxel of typical 2mm isotropic linear dimensions. The minimum value for *k* is four studies because at least that many are needed to define a volume, and therefore density, in three dimensions. However, *k* is an unknown parameter in ABC and must be selected based on two constraints: 1) the number must be small because the density estimate is only valid for small volumes to meet anatomical constraints such as the thin cortical ribbon, and 2) the p-values resulting from the density estimate must be uniformly distributed for random coordinates for statistical thresholding to work. These requirements are considered in the random coordinate experiment section.

Given a relevant tissue volume, such as the GM volume *V_gm_*, the probability of a coordinate falling within *dV_i_* if placed uniformly at random within the volume is *dV_i_/V_gm_*. If study *j* reports *C_j_* coordinates the probability of at least one of them falling within volume *dV_i_* is

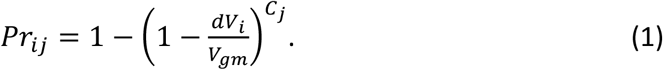

The p-value for coordinate *i* is the probability of *s≥k* coordinates from different studies falling in volume *dV_i_* assuming they are uniformly distributed in *V_gm_*, which for *N* studies is

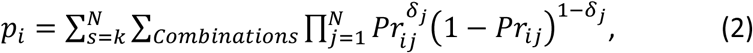

where *δ_j_* is either 0 or 1 and the sum over unique combinations includes all with

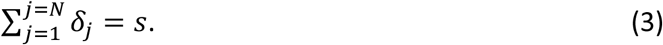

Combinations are found using Heap’s algorithm (Heap 1963). One implementation note is that equation (2) is generally more efficiently computed by summing *s* from 0 to *k*-1 and subtracting from 1.

### Forming clusters of high study density

The purpose of clustering in CBMA is to identify isolated anatomical regions that are associated with the hypothesised effect considering evidence from all studies. Most CBMA algorithms form clusters from spatially separated islands of voxels where the test statistic is greater than some threshold but this does require extrapolation of the coordinates, which is usually done using a fixed empirical smoothing kernel. In ABC the clustering is not voxel-wise but coordinate-wise and involves only coordinates that survive statistical thresholding. The approach is based on mean shift clustering (Fukunaga and Hostetler 1975), which shifts coordinate *i* in the direction of the weighted mean of other nearby coordinates from other studies. Iteratively performing this mean-shift operation drives coordinates towards isolated cluster centres. The process is complete when the shifted coordinate and the mean coincide. To proceed a kernel is required so that the shift towards the mean can be estimated, and in ABC the kernel takes the form of

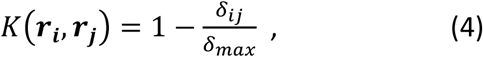

where *δ_j_* is the distance between coordinates ***r_i_*** and ***r_j_*** (*δ_ij_* =|***r_i_ – r_j_***|) and *δ_max_* is the largest distance parameter. Importantly the kernel width is not fixed empirically, instead it is deduced from an optimisation process; see the section on *choosing the kernel width.* The kernel is zero for *δ_ij_* > *δ_max_* to avoid influence from coordinates that are separated by large distances.

The shift vector for coordinate *i* involves a kernel weighted sum over all nearest coordinates from studies other than the study to which *i* belongs.

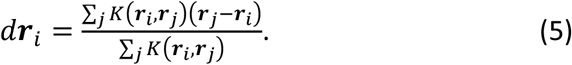

The algorithm iterates the calculation of distance between coordinates and application of the shift (equation (5)) to update the coordinates. The number of clusters detected by this method does not need to be specified a-priori. Of the clusters formed, each study may only contribute at most a single coordinate. If multiple coordinates from the same study apparently contribute, only the most significant (smallest p-value) is recruited into the cluster. A further requirement is that the number of studies contributing a coordinate to the cluster exceeds a minimum, which is determined as part of the principled statistical thresholding; see thresholding the coordinate p-values.

#### Choosing the kernel width

The mean shift clustering algorithm requires the specification of parameter *δ_max_*, which is somewhat analogous to the FWHM parameter used in other CBMA methods. However, in ABC it is automatically estimated specifically for the studies being analysed rather than being empirically specified once for all analyses. Only the coordinates that have survived statistical thresholding in ABC are considered for clustering, which makes the task simpler because the non-significant coordinates that fall sparsely between the dense clusters are not. The chosen value for *δ_max_* is that which maximises the number of the significant coordinates that are clustered. If *δ_max_* is set too low, then clusters are formed by too few studies to be valid, while if set too large clusters can merge reducing the number of clustered coordinates because studies may only contribute a single coordinate to any cluster. The search for the optimal value is performed by systematically searching between reasonable range of 3mm to 30mm in 0.1mm steps; this range can be widened if no optima are found.

### Thresholding the coordinate p-values

Principled statistical thresholding is important because the analyst must be able to specify some acceptable expected error rate. Other CBMA methods use fixed p-value thresholds, FDR, or FWE, and can be cluster-wise or voxel-wise. Fixed p-value thresholds offer no principled control over the false positives and are not recommended (Bennett et al. 2009) because error rates are determined by the data rather than being specified as appropriate by the analyst. Voxel-wise FDR allows the analyst to control the expected number of voxels falsely rejected under the null hypothesis as a proportion of total rejections, but these falsely rejected voxels can form spurious clusters that become part of the results and conclusions; because of this FDR is no longer the recommended threshold scheme for the popular ALE method (Eickhoff et al. 2016). Family wise error allows the analyst to control the proportion of analyses of null data that would produce significant results. Cluster-wise FWE prioritises larger clusters over smaller clusters, where the studies may report coordinates more densely, despite the latter potentially suggesting tighter agreement across studies. Perhaps the biggest limitation of the various methods employed is that non can easily be related to the problem of detecting replication of effects across studies.

Because ABC computes p-values coordinate-wise rather than voxel-wise, it is possible to relate the statistical threshold directly to the replicability of a result making the consequence of choosing a particular threshold easy to interpret. The method is somewhat analogous to FDR, but instead of controlling for the proportion (often 5%) of the rejections that are expected under the null hypothesis, it controls for the number of rejections expected under the null. In keeping with the primary idea of CBMA, the analyst chooses how replicable a result must be to be of interest in their expert opinion. This is done by specifying the proportion *(α)* of the studies that must contribute to a cluster for it to be considered indicative of a replicable effect, and from this a threshold is deduced. For number of studies *N,* and the total number of coordinates *N_c_*, the threshold (*p_thresh_*) is the largest p-value computed using equation (2) to obey the inequality

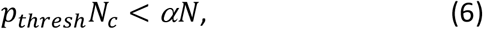

which constrains the expected number of coordinates falsely declared significant (under the null) to be fewer than the number of studies needed to indicate replicability in the analyst’s opinion. The threshold is generally more stringent than FDR, but FDR is also employed as an upper limit to provide FWE control under the null hypothesis (Benjamini and Hochberg 1995) so that when studies show no evidence of spatially consistent effects the algorithm is unlikely to return significant results. An implementation note is that a minimum of *k* studies must contribute to a cluster (*αN≥k*), where *k* is the number of studies used in calculating the p-values, and *k* must be at least 4 to define a volume in three dimensions.

A feature of this method is that the analyst must consider the proportion *α* carefully because there is a trade-off between the desire to detect more clusters and the need for those clusters to be significant. For example, if the analyst requires only a small proportion in the hope of finding more valid clusters, then the p-value threshold becomes more stringent as is clear from equation (6). Another important feature is that it considers studies that report no coordinates. An analyst requiring 25% of studies to contribute a coordinate to a cluster for it to be of interest does so regardless of those studies reporting no coordinates; consider that it is easier to achieve 25% of studies contributing to a cluster if coordinates are reported by all studies compared to only half of studies. Consequently, studies that report no significant coordinates, which is suggestive of no detectable hypothesis related effects, impose themselves on the analysis by making valid clustering more difficult to achieve.

### Analysing multiple similar hypotheses

Simultaneous analysis of related hypotheses is also possible using ABC. An example considered in this report is meta-analysis of fMRI studies of painful stimulus. There are several common ways of applying a painful stimulus, for examples electrical, mechanical, and thermal, and the stimulus factor might introduce some heterogeneity. If the studies were naively combined results specific to stimulus may be lost among the more significant common-to-all results. In ABC the significance of the coordinates is considered per stimulus type before they are combined for statistical thresholding and clustering. In this way the p-values for thermal stimulus coordinates, for example, are independent of the mechanical and electrical stimulus coordinates.

### Including multiple within-subject analyses

Typically, a study may report tables of coordinates from multiple analyses on the same subjects, for example painful stimulation by heat and by mechanical pressure; it is also possible that these analyses are reported in multiple papers. These should not be considered independent evidence of effect, so it is recommended that coordinates are arranged by subject group to form one independent study (Turkeltaub et al. 2012). However, this can increase the number of coordinates without respective increase in independent effects; consider that two within-subject group analyses producing perfectly correlated results would double the number of coordinates but not evidence of effect. In ABC this adversely impacts sensitivity by artifactually increasing the probabilities from equation (1). This issue also impacts other methods where coordinates are assumed independently and identically distributed under the null hypothesis such as ALE (Eickhoff et al. 2012); the multilevel kernel density analysis (MKDA) algorithm attempts to address this using a concept of empirical within-subject group coordinate clusters (Wager et al. 2007).

In ABC, two approaches have been implemented to reduce the impact of multiple correlated within-subject group analyses. Firstly, ABC automatically removes any within-subject group coordinate duplicates; such duplicates cannot be considered independent evidence of the same effect but equally should not reduce sensitivity to real effects. Secondly, ABC can be used to analyse multiple within-subject analyses that differ by a factor, for example painful stimulus (thermal, electrical, mechanical). Analysis is performed independently on each level of the factor so that they are unaffected by correlated resulst. Another possible approach is to include only the most relevant table from each study, which will also minimise heterogeneity, but might miss some relevant result. However, multiple within-subject group analyses could be included if they do not correlate strongly, and ABC automatically saves the correlation; correlation (Pearson’s) is of the spatial pattern of activation computed by fitting a Gaussian of FWHM=10mm to each coordinate (similarly to ALE) from two within-subject group analyses. For highly correlated results a sensitivity analysis may be advisable, performed by including only one of the correlated results, randomly selected over multiple analyses, per subject-group.

### Experiments

In this report the ABC concept is demonstrated using simulated and real data. In each example the grey matter volume, required for the probability model, is considered to be 780ml, which is the mean of the reported average grey matter volume in females and males (Lüders et al. 2002).

Two examples are considered using coordinates from fMRI studies employing painful stimulation. In example 1, 22 independent studies applying mechanically induced pain are analysed using both coordinates and reported statistical effects; these data have been used and provided previously (C. R. Tench et al. 2020). In the second example studies of painful stimulation by three different mechanisms are considered independently for coordinate inference before being merged for clustering using ABC: 28 mechanical pain stimulation studies, 29 electrical pain stimulation, and 83 thermal pain stimulation. These data files have also been used, and provided, previously (Tanasescu et al. 2016).

#### Experiments with random coordinates

It is important that random coordinates, representing the null distribution of ABC, are unlikely to produce significant clusters. Type 1 errors are controlled such that the expected number of false positive coordinates is fewer than necessary to form a valid cluster by imposing the inequality in equation (6) and by an upper limit imposed by FDR, but for this to work correctly the p-values must be uniformly distributed for random coordinates. Coordinates from example 1 are randomised uniformly into a grey matter mask and ABC performed 100 times. For each iteration, the number of clusters formed are counted. Across all 100 iterations the distribution of the p-values is also recorded. This procedure is performed using the *k=4* and *k=5* nearest studies when estimating study density. The number of random experiments producing clusters is reported and the cumulative p-value distribution plotted.

#### Painful stimulus fMRI study including statistical effect size

Analysis of brain coordinates is performed on the 22 independent studies of mechanically induced pain and presented for comparison with the same data were analysed using the ALE algorithm (Eickhoff et al. 2012) (GingerALE version 3.0.2, cluster-wise thresholding: FWE=0.05, cluster forming threshold p=0.001). The resulting p-values computed using equation (2) are depicted as Z-scores, and the resulting clusters are depicted as coloured regions of interest (ROIs). In this analysis 5 (~23%) studies are required to make a valid cluster.

#### Electrical, mechanical, and thermal painful stimulus

ABC is performed on these independent fMRI studies. The p-values are depicted as Z-scores and clusters as coloured ROIs. In this analysis 5 (~3.6%) studies are required to make a valid cluster.

## Results

### Random experiments

Using *k*=4 studies to estimate the study coordinate density resulted in 20 out of 100 analyses using random coordinates producing significant clusters. This means that studies with no better spatial agreement than random coordinates can still form significant results around 1 in 5 times. Methods such as ALE use FWE such that with random coordinates would produce significant results only 1 in 20 times (for default FWE of 0.05). The rate of 1 in 5 positive results under random conditions is undesirably high when the purpose of CBMA is to capture the results systematically repeated across the studies. However, when *k=5* studies are used to estimate the study coordinate density none of the 100 iterations produced significant clusters. The explanation for this is that the p-values of random coordinates are not uniformly distributed, as required under the null hypothesis, when the study density is estimated using *k=*4 studies (figure 1, dashed line). Consequently, the principled error rate control will not operate as expected because the number of coordinates with small p-values is greater than expected. However, using *k=*5 studies to estimate study density produces p-values that are closer to uniformly distributed (figure 1, solid line) for random coordinates as required.

**Figure 1.**
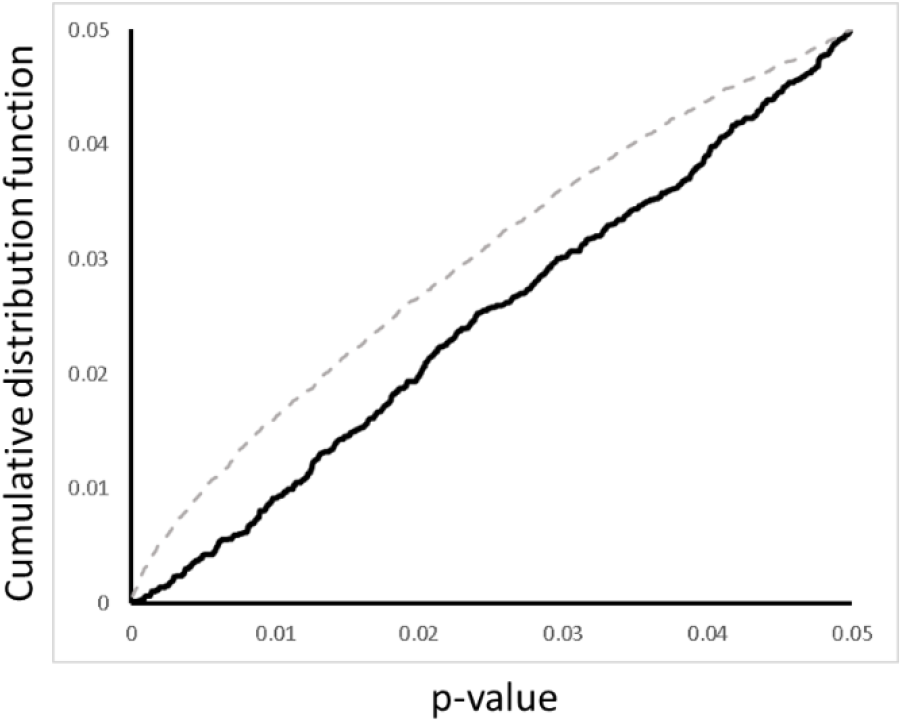
The cumulative distribution of random coordinate p-values computed using *k*=4 (dashed line) and *k*=5 (black line) studies to estimate the study density. For *k*=5 the p-values are uniformly distributed as required.

### Painful stimulus fMRI study

The coordinate significance and clusters found on estimating 22 independent fMRI studies of mechanically induced pain are depicted in figure 2. For comparison, the results from the ALE algorithm are also provided (top). Both algorithms produce a similar result, but some differences are evident.

**Figure 2.**
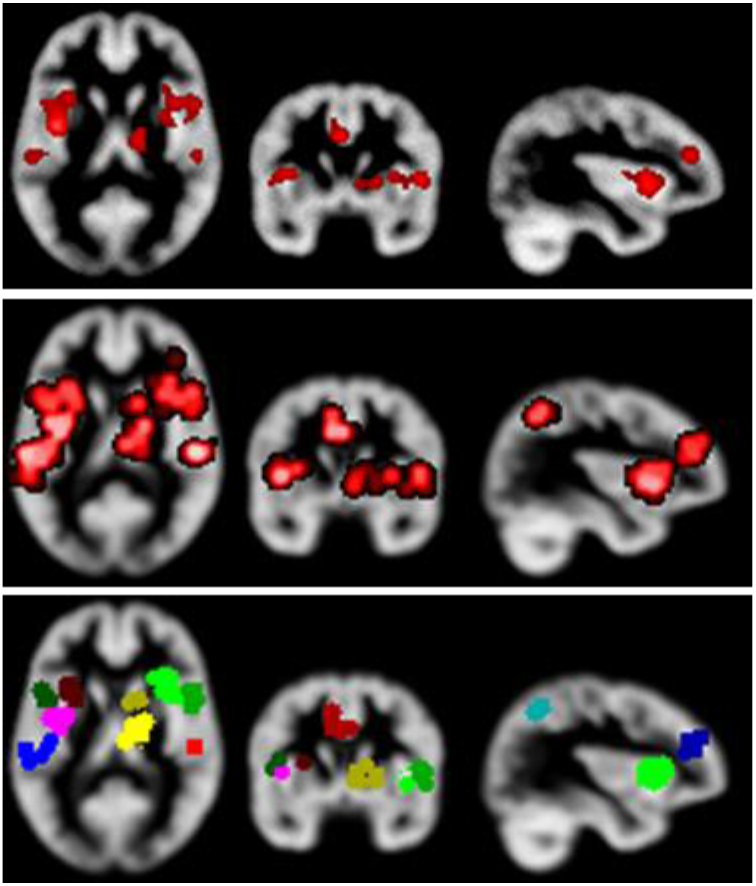
ABC of 22 fMRI studies of mechanically induced pain. The top image depicts the results produced using the ALE algorithm (cluster-based thresholding, FWE=0.05, cluster forming threshold p=0.001). The middle image shows the significance (displayed as Z scores) deduced using the ABC algorithm; a small amount of smoothing has been applied to improve visualisation. The clustered significant coordinates produced using ABC are shown at the bottom.

**Figure 3.**
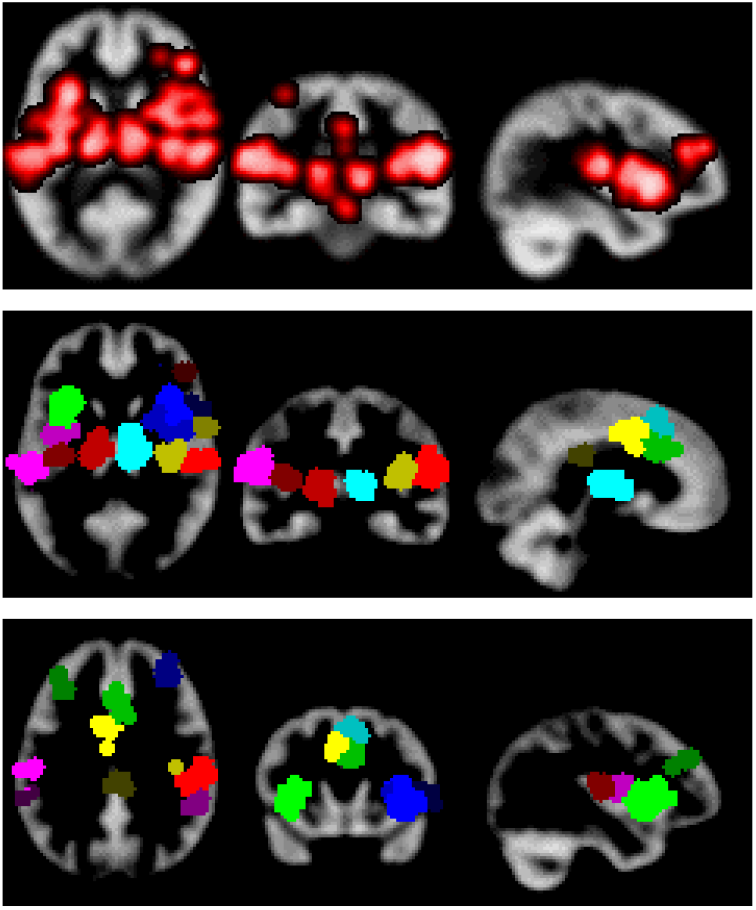
ABC of three different modes of painful stimulation; thermal, mechanical, and electrical. The top image shows the significance of the coordinates overlaid on a grey matter image; a small amount of smoothing has been applied to aid visualisation. The middle and bottom images depict two views of the resulting clusters.

ABC was performed on the 140 fMRI studies of painful stimulus separated by stimulus modality (thermal, mechanical, electrical) and results are shown in figure 5. The results reveal multiple clusters placed with considerable hemispheric symmetry. Note no comparison with ALE results is provided as no similar analysis is available.

## Discussion

Here a method of performing a meta-analysis of functional MRI or voxel-based morphometry studies has been presented. The aim of ABC is to provide analysis where direct influence of the algorithm on the results is relatively simple to understand. Just as with other CBMA algorithms ABC can help further understanding of brain function by providing clear summaries of results from multiple studies and can even be used to test hypotheses if results can be predicted independently before performing the study.

Amongst the various CBMA algorithms the closest to ABC is parametric voxel-based metaanalysis (PVM) (Sergi G. Costafreda et al. 2009). A model of study density, similar to that in ABC, is used voxel-wise before thresholding to control the FDR. ABC offers some potential advantages over PVM: 1) because of the computational demands of computing the voxelwise p-values the PVM method uses approximation, 2) PVM uses empirical fixed kernel width, which has been shown to cause clusters to grow in size with increasing number of studies (Eickhoff et al. 2016; Christopher R Tench et al. 2014) and therefore produces results that are not convergent and can paradoxically increase the proportion of false positive results (Christopher R Tench et al. 2014) as the expanding clusters recruit the noisy coordinates falling between clusters, 3) the use of voxel-wise FDR is known to cause issues in CBMA because the expected proportion of results that are falsely declared significant (the false discovery rate) can be sufficient to produce spurious clusters (Eickhoff et al. 2016), and 4) studies differing by a factor, and within the same subject group, can be analysed with p-values computed independently per factor level avoiding problems with correlated results.

Using a model-based approach avoids the need to randomise coordinates into an empirical preselected image space, instead needing only the volume of interest such as the grey matter volume. An advantage of the coordinate-wise model-based approach is computational efficiency, with typically sized analyses involving a few tens of studies taking just seconds. By using the local density of studies, the requirement for a fixed empirical smoothing kernel is also avoided. A primary aim of CBMA is to filter those results that are study specific, perhaps due to use of uncorrected voxel-wise testing for example, leaving those that appear consistent across study, which are more likely to be hypothesis specific. A convenient and interpretable threshold must be applied to achieve this filtering, and ABC thresholds the model-based p-values such that the expected number of coordinates falsely declared significant is less than a user selected proportion of studies. The expert analyst preselects this proportion by consideration of the minimum to consider the result replicable.

In part it is the disparity of results available from the multiple neuroimaging analysis packages that motivate the development of CBMA. Ironically, there are also multiple algorithms for performing CBMA, each with different empirical features and producing a range of results (Ferreira and Busatto 2010). Coordinate based meta-analyses can therefore be improved by providing a protocol in which the analyst specifies and justifies the methodology a-priori (Tahmasian et al. 2019), removing the option to choose the algorithm or settings giving the best result, which may be biased by analyst opinion. ABC has been developed specifically to be easy to understand and think about prospectively, which has been achieved by eliminating some empirical assumptions. Requirements of performing and reporting ABC analysis are similar to those of meta-analysis and CBMA (Müller et al. 2018). CBMA assumes that studies are independent. It is important that multiple experiments on the same subjects are not considered independent as this will produce a known form of bias common to meta-analysis, and consequently reduce the quality of evidence. It is also important to provide the data analysed along with any publication; typically, multiple experiments are reported per study, and it may be difficult to know which experiments have been included, and therefore to interpret the results or reproduce the analysis. Provision of data in any meta-analysis is a PRISMA (Preferred Reporting Items for Systematic Reviews and Meta-Analyses) requirement, and only involves inclusion of text files.

## Summary and conclusions

Meta-analysis is considered very high-level evidence. Its importance in neuroimaging is in identifying those published results that are replicable across multiple studies where potential for false positive or study specific effects is high. There are many algorithms, but each produces different results that are always conditional on the empirical assumptions used and options chosen. It is therefore important to plan the study in detail to avoid making analysis decisions after seeing the results, which might be a source of bias. ABC uses minimal empirical assumptions and an easy to interpret principled method of error control, making CBMA using ABC simple to plan and understand.

## Funding

No specific funding is associate with this research

RT received support in part from MRC (CARP MR/T024402/1)

## Conflicts of interest/Competing interests

There are no conflicts of interest

## Availability of data and material

Data used in this paper have been made available elsewhere

## Code availability

Software to perform analysis is made freely available https://www.nottingham.ac.uk/research/groups/clinicalneurology/neuroi.aspx

## Ethics approval

No ethics required for this methods paper

## Notes

### Competing Interest Statement

The authors have declared no competing interest.

### Summary of Updates

This revision does not represent a change in the methodology. It renames the algorithm to emphasize its purpose, and describes the advantages it offers.

## References

Albajes-Eizagirre, A., Solanes, A., Vieta, E., & Radua, J. (2018). Voxel-based meta-analysis via permutation of subject images (PSI): Theory and implementation for SDM. Neuroimage, 186, 174–184. https://doi.org/10.1016/j.neuroimage.2018.10.077

Benjamini, Y., & Hochberg, Y. (1995). Controlling the False Discovery Rate: A Practical and Powerful Approach to Multiple Testing. Journal of the Royal Statistical Society. Series B (Methodological), 57(1), 289–300. https://doi.org/10.2307/2346101

Bennett, C. M., Wolford, G. L., & Miller, M. B. (2009). The principled control of false positives in neuroimaging. Soc Cogn Affect Neurosci, 4(4), 417–22. https://doi.org/10.1093/scan/nsp053

Costafreda, S. G. (2012). Parametric coordinate-based meta-analysis: valid effect size meta-analysis of studies with differing statistical thresholds. J Neurosci Methods, 210(2), 291–300. https://doi.org/10.1016/j.jneumeth.2012.07.016

Costafreda, Sergi G., David, A. S., & Brammer, M. J. (2009). A parametric approach to voxel-based meta-analysis. NeuroImage, 46(1), 115–122. https://doi.org/10.1016/j.neuroimage.2009.01.031

Eickhoff, S. B., Bzdok, D., Laird, A. R., Kurth, F., & Fox, P. T. (2012). Activation likelihood estimation meta-analysis revisited. Neuroimage, 59(3), 2349–61. https://doi.org/S1053-8119(11)01062-7 [pii] 10.1016/j.neuroimage.2011.09.017

Eickhoff, S. B., Laird, A. R., Grefkes, C., Wang, L. E., Zilles, K., & Fox, P. T. (2009). Coordinate-based activation likelihood estimation meta-analysis of neuroimaging data: a randomeffects approach based on empirical estimates of spatial uncertainty. Hum Brain Mapp, 30(9), 2907–26. https://doi.org/10.1002/hbm.20718

Eickhoff, S. B., Nichols, T. E., Laird, A. R., Hoffstaedter, F., Amunts, K., Fox, P. T., et al. (2016). Behavior, sensitivity, and power of activation likelihood estimation characterized by massive empirical simulation. Neuroimage, 137, 70–85. https://doi.org/10.1016/j.neuroimage.2016.04.072

Ferreira, L. K., & Busatto, G. F. (2010). Heterogeneity of coordinate-based meta-analyses of neuroimaging data: an example from studies in OCD. The British Journal of Psychiatry, 197(1), 76–77. https://doi.org/10.1192/bjp.197.1.76a

Fukunaga, K., & Hostetler, L. (1975). The estimation of the gradient of a density function, with applications in pattern recognition. IEEE Transactions on Information Theory, 21(1), 32–40. Presented at the IEEE Transactions on Information Theory. https://doi.org/10.1109/TIT.1975.1055330

Heap, B. R. (1963). Permutations by Interchanges. The Computer Journal, 6(3), 293–298. https://doi.org/10.1093/comjnl/6.3.293

Kiefer, J. (1953). Sequential minimax search for a maximum. Proceedings of the American Mathematical Society, 4(3), 502–506. https://doi.org/10.1090/S0002-9939-1953-0055639-3

Laird, A. R., Fox, P. M., Price, C. J., Glahn, D. C., Uecker, A. M., Lancaster, J. L., et al. (2005). ALE meta-analysis: controlling the false discovery rate and performing statistical contrasts. Hum Brain Mapp, 25(1), 155–64. https://doi.org/10.1002/hbm.20136

Li, C., Liu, W., Guo, F., Wang, X., Kang, X., Xu, Y., et al. (2020). Voxel-based morphometry results in first-episode schizophrenia: a comparison of publicly available software packages. Brain Imaging and Behavior, 14(6), 2224–2231. https://doi.org/10.1007/s11682-019-00172-x

Lüders, E., Steinmetz, H., & Jäncke, L. (2002). Brain size and grey matter volume in the healthy human brain. Neuroreport, 13(17), 2371–2374. https://doi.org/10.1097/01.wnr.0000049603.85580.da

Müller, V. I., Cieslik, E. C., Laird, A. R., Fox, P. T., Radua, J., Mataix-Cols, D., et al. (2018). Ten simple rules for neuroimaging meta-analysis. Neuroscience & Biobehavioral Reviews, 84, 151–161.

Popescu, V., Schoonheim, M. M., Versteeg, A., Chaturvedi, N., Jonker, M., Xavier de Menezes, R., et al. (2016). Grey Matter Atrophy in Multiple Sclerosis: Clinical Interpretation Depends on Choice of Analysis Method. PLoS One, 11(1), e0143942. https://doi.org/10.1371/journal.pone.0143942

Radua, J., Mataix-Cols, D., Phillips, M. L., El-Hage, W., Kronhaus, D. M., Cardoner, N., & Surguladze, S. (2012). A new meta-analytic method for neuroimaging studies that combines reported peak coordinates and statistical parametric maps. Eur Psychiatry, 27(8), 605–11. https://doi.org/S0924-9338(11)00073-3 [pii] 10.1016/j.eurpsy.2011.04.001

Tahmasian, M., Sepehry, A. A., Samea, F., Khodadadifar, T., Soltaninejad, Z., Javaheripour, N., et al. (2019). Practical recommendations to conduct a neuroimaging meta-analysis for neuropsychiatric disorders. Human Brain Mapping, 40(17), 5142–5154. https://doi.org/10.1002/hbm.24746

Talairach, J., & Tournoux, P. (1988). Co-planar stereotaxic atlas of the human brain. New York: Thieme.

Tanasescu, R., Cottam, W. J., Condon, L., Tench, C. R., & Auer, D. P. (2016). Functional reorganisation in chronic pain and neural correlates of pain sensitisation: a coordinate based meta-analysis of 266 cutaneous pain fMRI studies. Neuroscience & Biobehavioral Reviews, 68, 120–133.

Tench, C. R., Tanasescu, R., Constantinescu, C. S., Auer, D. P., & Cottam, W. J. (2017). Coordinate based random effect size meta-analysis of neuroimaging studies. Neuroimage, 153, 293–306. https://doi.org/10.1016/j.neuroimage.2017.04.002

Tench, C. R., Tanasescu, R., Constantinescu, C. S., Cottam, W. J., & Auer, D. P. (2020). Coordinate based meta-analysis of networks in neuroimaging studies. NeuroImage, 205, 116259. https://doi.org/10.1016/j.neuroimage.2019.116259

Tench, Christopher R, Tanasescu, R., Auer, D. P., Cottam, W. J., & Constantinescu, C. S. (2014). Coordinate based meta-analysis of functional neuroimaging data using activation likelihood estimation; full width half max and group comparisons. PloS one, 9(9), e106735.

Turkeltaub, P. E., Eden, G. F., Jones, K. M., & Zeffiro, T. A. (2002). Meta-analysis of the functional neuroanatomy of single-word reading: method and validation. Neuroimage, 16(3 Pt 1), 765–80. https://doi.org/S1053811902911316 [pii]

Turkeltaub, P. E., Eickhoff, S. B., Laird, A. R., Fox, M., Wiener, M., & Fox, P. (2012). Minimizing within-experiment and within-group effects in Activation Likelihood Estimation meta-analyses. Hum Brain Mapp, 33(1), 1–13. https://doi.org/10.1002/hbm.21186

Wager, T. D., Lindquist, M., & Kaplan, L. (2007). Meta-analysis of functional neuroimaging data: current and future directions. Soc Cogn Affect Neurosci, 2(2), 150–8. https://doi.org/10.1093/scan/nsm015

Wager, T. D., Phan, K. L., Liberzon, I., & Taylor, S. F. (2003). Valence, gender, and lateralization of functional brain anatomy in emotion: a meta-analysis of findings from neuroimaging. Neuroimage, 19(3), 513–31.

